# High genetic barrier to escape from human polyclonal SARS-CoV-2 neutralizing antibodies

**DOI:** 10.1101/2021.08.06.455491

**Authors:** Fabian Schmidt, Yiska Weisblum, Magdalena Rutkowska, Daniel Poston, Justin Da Silva, Fengwen Zhang, Eva Bednarski, Alice Cho, Dennis J. Schaefer-Babajew, Christian Gaebler, Marina Caskey, Michel C. Nussenzweig, Theodora Hatziioannou, Paul D. Bieniasz

**Affiliations:** Laboratory of Retrovirology, The Rockefeller University, New York, NY 10065, USA; Laboratory of Molecular Immunology, The Rockefeller University, New York, NY 10065, USA; Howard Hughes Medical Institute

## Abstract

The number and variability of the neutralizing epitopes targeted by polyclonal antibodies in SARS-CoV-2 convalescent and vaccinated individuals are key determinants of neutralization breadth and, consequently, the genetic barrier to viral escape. Using chimeric viruses and antibody-selected viral mutants, we show that multiple neutralizing epitopes, within and outside the viral receptor binding domain (RBD), are variably targeted by polyclonal plasma antibodies and coincide with sequences that are enriched for diversity in natural SARS-CoV-2 populations. By combining plasma-selected spike substitutions, we generated synthetic ‘polymutant’ spike proteins that resisted polyclonal antibody neutralization to a similar degree as currently circulating variants of concern (VOC). Importantly, by aggregating VOC-associated and plasma-selected spike substitutions into a single polymutant spike protein, we show that 20 naturally occurring mutations in SARS-CoV-2 spike are sufficient to confer near-complete resistance to the polyclonal neutralizing antibodies generated by convalescents and mRNA vaccine recipients. Strikingly however, plasma from individuals who had been infected and subsequently received mRNA vaccination, neutralized this highly resistant SARS-CoV-2 polymutant, and also neutralized diverse sarbecoviruses. Thus, optimally elicited human polyclonal antibodies against SARS-CoV-2 should be resilient to substantial future SARS-CoV-2 variation and may confer protection against future sarbecovirus pandemics.

---

Neutralizing antibodies elicited by prior infection or by vaccination likely represent a key component of protective immunity against SARS-CoV-2. Antibodies targeting the receptor binding domain (RBD) of the spike protein are thought to dominate the neutralizing activity of convalescent or vaccine recipient plasma^1^, and include the most potent neutralizing antibodies cloned from convalescent individuals^2–6^. However, additional SARS-CoV-2 neutralizing antibody targets include the N-terminal domain (NTD) and the fusion machinery^3,5,7–9^, and the full spectrum of epitopes targeted by neutralizing antibodies in convalescent or vaccine recipient plasma has not been defined. Circulating SARS-CoV-2 variants of concern (VOC) or variants of interest (VOI) encode spike amino acid substitutions^10–13^, some of which confer resistance individual human monoclonal antibodies but have variable, typically modest, effects on neutralization by polyclonal plasma antibodies^1,4,13–17^. Mutated sites in VOCs include those in the RBD, NTD and elsewhere, but the numbers and locations of spike substitutions required for SARS-CoV-2 to evade the polyclonal neutralizing antibodies encountered in vaccine recipients or convalescents is unknown, and is a crucial determinant of population immunity.

## Targets of polyclonal neutralizing antibodies

Exploiting the fact that SARS-CoV is poorly neutralized by SARS-CoV-2 convalescent plasma, we compared the sensitivity of HIV-1 pseudotypes bearing parental and chimeric spike proteins in which RBD sequences were exchanged (SARS-CoV-2(1-RBD) and SARS-CoV(2-RBD), Fig. 1a, Extended data Fig. 1) to neutralization by plasma from 26 individuals from a previously described Rockefeller University COVID19 convalescent cohort^18^. The plasma samples were obtained at an average of 1.3 months after infection and were selected for high SARS-CoV-2 neutralization titers (RU27 plasma panel). Compared to the SARS-CoV-2 pseudotype, the SARS-CoV-2(1-RBD) pseudotype was less sensitive to neutralization by 21/26 plasmas (median difference = 1.8-fold, range 0.5 to 9.8-fold, p=0.0005, Fig. 1b), while SARS-CoV(2-RBD) pseudotype was more sensitive to all plasmas (median difference 8-fold, range 1.2 to 75.5 fold, p<0.0001, Fig 1c). Nevertheless, the neutralizing potency of some plasmas was hardly affected when the SARS-CoV-2 RBD was replaced by the SARS-CoV RBD, even though some of those plasmas were minimally active against SARS-CoV (e.g. RU9, RU10, RU11, RU15 Fig. 1b, c). Indeed, plasmas that poorly neutralized SARS-CoV potently neutralized chimeric spike pseudotypes with either RBD or the other spike domains from SARS-CoV-2 (Fig. 1b, c). For the overall plasma panel, neutralizing potency against SARS-CoV-2 pseudotypes did not correlate with potency against SARS-CoV pseudotypes or SARS-CoV-2(1-RBD) (Fig 1d,e), but did correlate with potency against SARS-CoV(2-RBD) (Fig 1f). Although altered RBD conformation in chimeric spike proteins might affect neutralization, these data indicate that while the RBD constitutes a dominant neutralizing target, substantial plasma neutralizing activity is also directed against non-RBD epitopes.

**Fig. 1.**
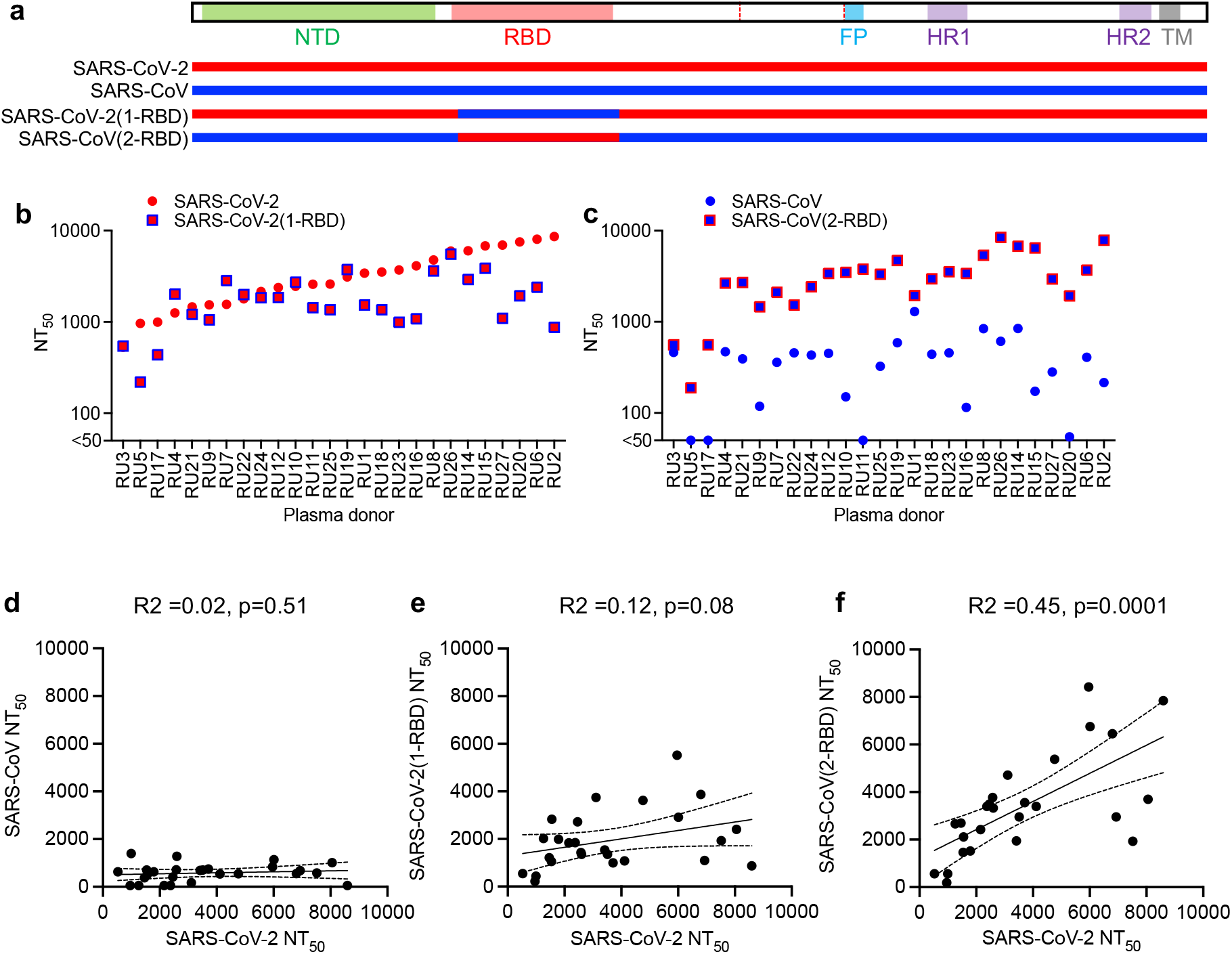
Neutralizing antibodies in SARS-CoV-2 convalescent plasma targeting both RBD and non-RBD determinants. (**a**) Design of RBD-exchanged chimeric spike proteins. (**b,c**) Fifty percent neutralization titers (NT_50_) for 26 high titer convalescent plasmas (RU1-27) against pseudotyped HIV-1 virions bearing the indicated spike proteins. Median of two independent experiments is plotted. (**d-f**) Correlations of neutralizing potencies of the plasmas against the indicated pairs of spike proteins.

## Selection of spike mutations by convalescent polyclonal antibodies

To more precisely map the targets of polyclonal neutralizing antibodies in convalescent individuals, we passaged a rVSV/SARS-CoV-2 chimeric virus^14,19^ in the presence of each of the RU27 plasmas for up to six passages. Importantly, rVSV/SARS-CoV-2 mimics the neutralization properties of SARS-CoV-2^14,19^ but obviates the safety concerns that would accompany such studies with authentic SARS-CoV-2. NGS sequencing indicated that rVSV/SARS-CoV-2 passage in 22 out of the 27 plasmas, led to the selection of spike mutations (Fig. 2a, Extended Data Fig 2, Table S1). For some plasmas, multiple mutations were selected at distinct but proximal sites in subpopulations, indicating a dominant neutralizing activity targeting a particular epitope. For other plasmas, selected mutations were enriched in multiple regions in the spike coding sequence suggesting co-dominant neutralizing activities. Six passages of the rVSV/SARS-CoV-2 chimeric virus without plasma enriched a small number of substitutions that were assumed to represent culture adaptation or fitness-enhancing mutations (e.g. T604I) but were distinct from the majority of mutations arising after rVSV/SARS-CoV-2 passage in plasma (Extended Data Fig. 2, Table S1). Cumulatively, the plasma-selected mutations were enriched in specific elements within NTD, RBD and other spike domains (Fig. 2a, Table S1). From the 27 plasma-passaged virus populations, 38 individual mutant viruses were isolated by plaque purification, each of which encoded one, two or three spike substitutions (Fig. 2b) that generally occurred at high frequency in the passaged viral populations (Table S1).

**Fig. 2.**
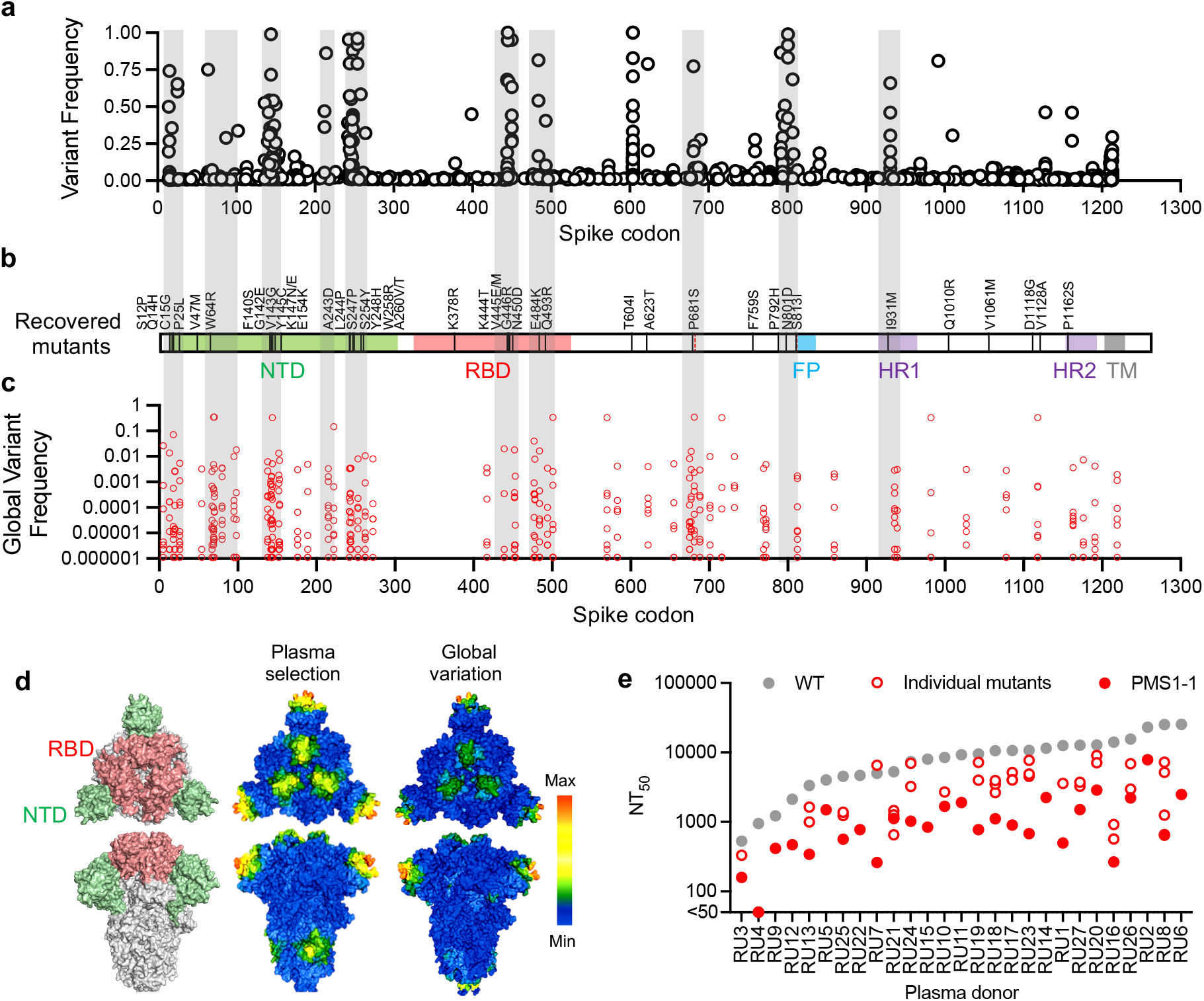
Selection of SARS-CoV-2 spike mutants by polyclonal antibodies. (**a**) Frequencies of amino acid substitutions at each codon of the SARS-CoV-2 spike protein in two independent rVSV-SARS-CoV-2 populations (1D7 and 2E1), determined by Illumina sequencing. Pooled results following selection with the RU27 plasma panel are displayed. (**b**) Locations of amino acid substitutions in 38 plaque purified rVSV/SARS-CoV-2 isolates obtained from rVSV/SARS-CoV-2 populations following passage in the RU27 plasmas. (**c**) Frequencies of naturally occurring amino acid substitutions (red circles) at each codon of the SARS-CoV-2 spike protein. Shaded gray bars in (**a-c**) indicate shared regions where variation is enriched. (**d**) Comparison of the averaged frequency of substitutions observed after passaging rVSV/SARS-CoV-2 with RU27 plasmas (center) and the frequency of sequence changes in natural populations (right), projected onto the SARS-CoV-2 spike structure (PDB 6VXX) with positions of the RBD and NTD domains indicated (right). The average frequency of substitutions in a 15 Å radius is represented using the color spectrum (scale = 0-20 center and 0-9 right). (**e**) Neutralization potency of RU27 plasmas against rVSV/SARS-CoV-2 encoding WT, individual selected mutants, or PMS1-1 spike proteins. Median of two independent determinations is plotted.

We compared the distribution of mutations selected by the RU27 plasma panel in cell culture with those occurring in naturally circulating SARS-CoV-2 populations (Fig. 2a–d). In both plasma-selected and naturally occurring spike sequences, substitutions were enriched in several elements that contribute to the NTD ‘supersite’ targeted by NTD-binding neutralizing antibodies^7,8^ (Fig. 2a–d). Similar plasma-selected and natural sequence variation was also evident in elements targeted by class 2 and class 3 RBD-binding neutralizing antibodies^20^. Mutations conferring resistance to class I RBD antibodies were not selected by plasma passage perhaps reflecting a lower than expected abundance of class I antibodies in this plasma panel (Fig. 2a–d). Other sites, including spike amino acids ~680-700 and ~930 exhibited variation in both plasma-passaged and natural variant datasets, but have not yet been demonstrated to be targeted by neutralizing antibodies. Nevertheless, the similarity in the distribution of natural and plasma-selected sequence variation within spike suggests that selection by neutralizing antibodies drives divergence in naturally circulating SARS-CoV-2 populations.

Of the 38 plaque-purified rVSV/SARS-CoV-2 mutants recovered following passage in RU27 plasmas, 34 exhibited varying degrees of reduced sensitivity to neutralization by the plasma that was used for its selection (median =3.1 fold reduced NT_50_, range 0.8 to 39.3 fold, Extended Data Fig. 3). Nevertheless, for 37/38 of the selected rVSV/SARS-CoV-2 mutants, the selecting plasma exhibited residual neutralizing activity. We aggregated 13 mutations from the plasma selected viruses based on their effects on plasma neutralization sensitivity (Extended data Fig. 3) and distribution throughout the spike protein, generating a single synthetic ‘polymutant’ spike (PMS) protein sequence, termed PMS1-1 (Extended Data Fig. 4a). Notably, an rVSV/SARS-CoV-2 derivative encoding these spike mutations (rVSV/SARS-CoV-2_PMS1-1_) exhibited resistance to neutralization by the RU27 plasma panel that was significantly greater in magnitude (p<0.0001) and consistency than the individual plasma selected mutants (median 8.0 -fold, range 2.7 to 52.9 fold, Fig. 2e, Extended Data Fig. 4b). Nevertheless, 26/27 of the RU27 plasmas retained residual neutralizing activity against rVSV/SARS-CoV-2_PMS1-1_ (Fig. 2e, Extended Data Fig. 4b). We conclude that some neutralizing epitopes are shared among the convalescent antibodies in high-titer plasmas, but neutralizing activity against SARS-CoV-2 is clearly polyclonal and heterogeneous among individuals with respect to epitope targets.

**Fig. 3.**
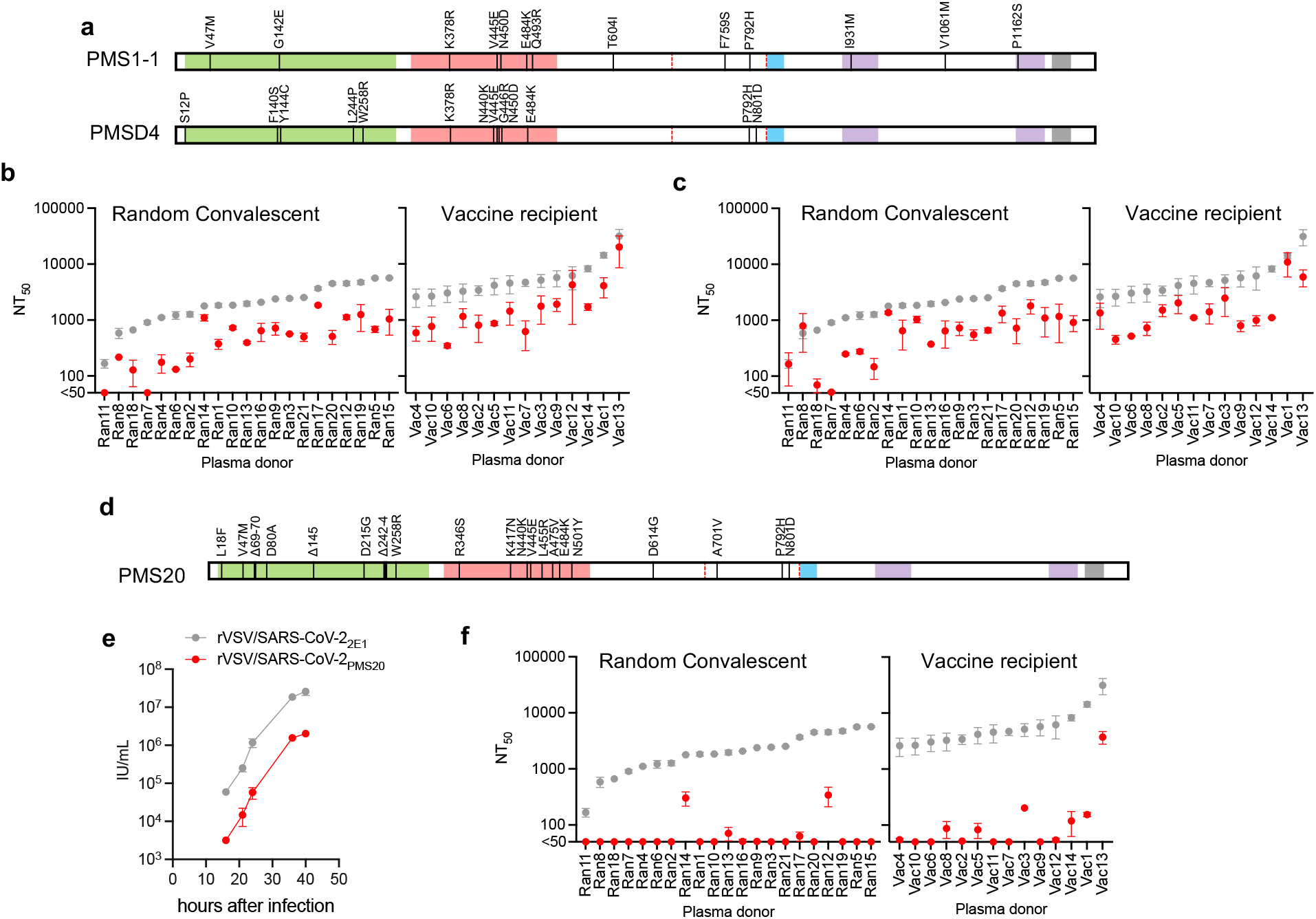
Neutralization resistance of polymutant SARS-CoV-2 spike proteins. (**a**) Design of a PMS1-1 and PMSD4 polymutant spike proteins with 13 plasma-selected spike mutations aggregated in each spike. (**b,c**) Comparative neutralization potency of randomly selected convalescent (Ran 1-21) and vaccine recipient (Vac1-14) plasmas, against Wuhan-hu-1 (grey symbols) and PMS1-1**(b)** or PMSD4 **(c)** (red symbols) SARS-CoV-2 HIV-1 pseudotypes. (**d**) Design of the PMS20 spike protein with 20 antibody-selected and VOC-associated mutations. (**e**) Replication of rVSV/SARS-CoV-2 chimeras encoding 2E1 (parental) or PMS20 spike proteins in 293T/ACE2cl.22 cells infected at a mutliplicity of 0.001 and 0.008 respectively. **(f)** Same as **b**,**c** but with the PMS20 spike protein. For **b**, **c** and **f** median and range of two independent determinations is plotted.

**Fig 4.**
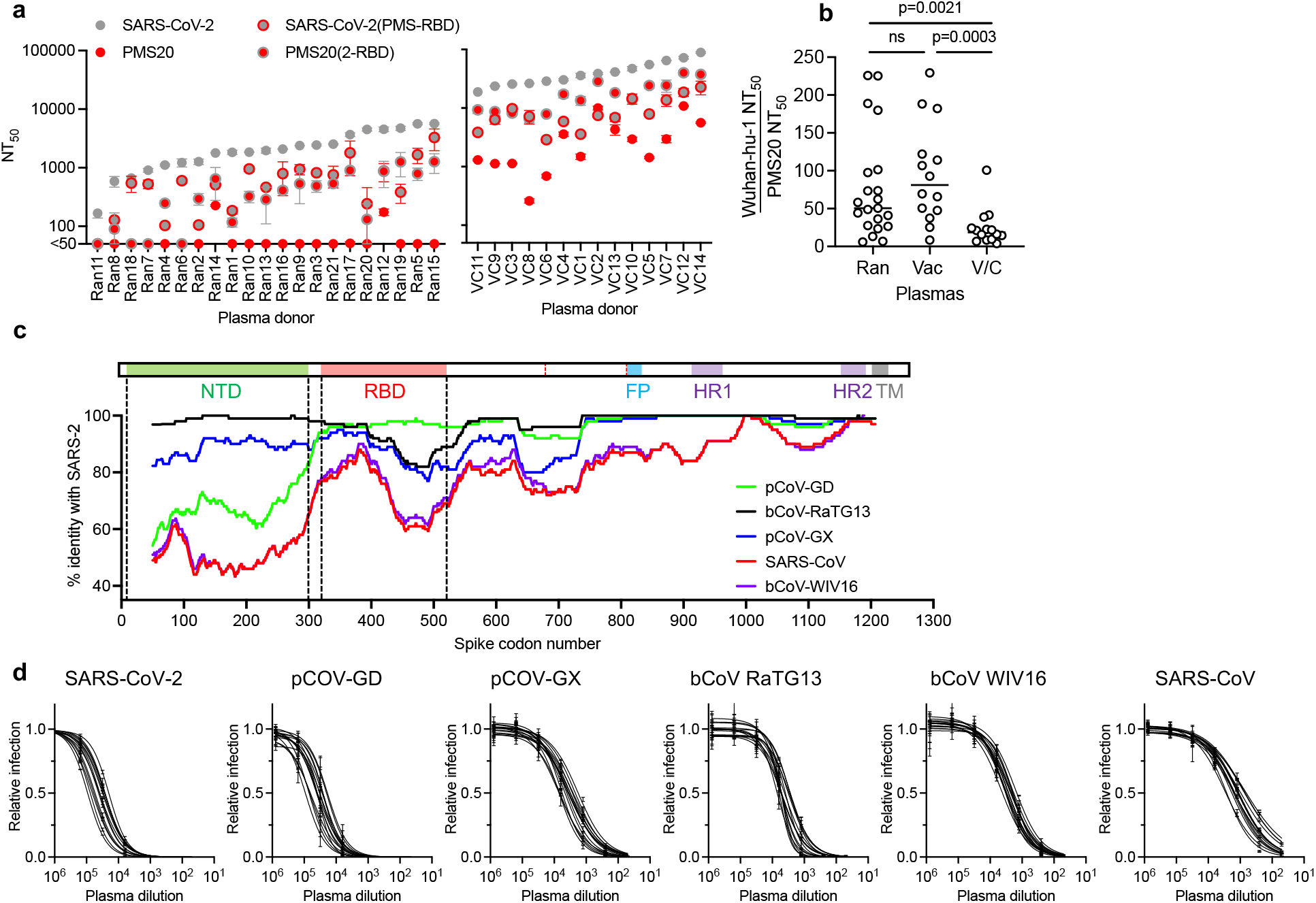
Neutralization breadth of polyclonal antibodies from vaccinated convalescents. (**a**) Comparative neutralization potency (NT_50_ values) of random convalescent (Ran1-21) and vaccinated convalescent (VC1-14) plasmas against HIV-1 pseudotypes bearing SARS-CoV-2, PMS20, and RBD-exchanged chimeric spike proteins. Median and range of two independent determinations is plotted. (**b**) Fold difference in NT_50_, comparing neutralization of HIV-1 pseudotypes bearing SARS-CoV-2 and PMS20 spike proteins by Ran1-21, Vac1-15 and VC1-14 plasmas. (**c**) Sequence diversity across sarbecovirus spike domains; SARS-CoV-2 and sarbecovirus spike sequences were aligned with Clustal and compared using Simplot; the percent identity relative to SARS-CoV2 was plotted within a rolling window of 100 amino acids. (**d**) Neutralization curves for 14 vaccinated convalescent plasmas and the indicated sarbecovirus HIV-1 pseudotypes.

## Synthetic polymutant and natural variant neutralization

We generated a panel of HIV-1 pseudotypes bearing the PMS1-1 spike protein, a second PMS protein with a different set of 13 mutations (selected based on similar criteria, PMSD4, Fig. 3a), or naturally occurring variants or relatives of SARS-CoV-2 spike (see Extended Data Fig. 1 for characterization of the pseudotypes). The panel included several SARS-CoV-2 VOC or VOI spike proteins (Extended Date Fig. 5a), and spike proteins from sarbecoviruses found in bats (bCoV-RaTG13), pangolins (pCoV-GD and pCOV-GX), and previously in humans (SARS-CoV), that exhibit varying degrees of sequence divergence from SARS-CoV-2^21–23^. To determine sensitivity/resistance to polyclonal SARS-CoV-2 antibodies, we employed an independent set of 21 randomly selected (Ran1-21) convalescent plasmas, and a set of 14 plasmas from mRNA vaccine recipients (Vac1-14), in addition to the RU27 high-titer plasma panel. The PMS spike proteins exhibited a degree of neutralization resistance that fell with the range of that exhibited by the four SARS-CoV-2 VOC/VOI and the four other sarbecoviruses. Specifically, PMS1-1 and PMSD4 exhibited neutralization resistance that was greater than B.1.1.7 and B.1.526, similar to P.1 and less than B.1.351.3 (Fig. 3b, c; Extended data Figs 5b and 6a, b). PMS1-1 and PMSD4 were more resistant to neutralization than pCoV-GD and bCoV-RaTG13 (Fig. 3b, c; Extended data Figs 5c and 6a, c), both of which contain a larger number of changes relative to SARS-CoV-2 than the PMS spike proteins. Conversely, the pCoV-GX and SARS-CoV pseudotypes were more resistant to SARS-CoV-2 convalescent and vaccine recipient plasma than PMS1-1 and PMSD4 (Extended data Figs 5c and 6c). Notably, like PMS1-1 and PMSD4, the B.1.351.3 VOC that encodes only nine spike mutations relative to SARS-CoV-2 Wuhan-hu-1, was more neutralization resistant than sarbecoviruses (pCoV-GD and bCoV-RaTG13) that contain a greater number of substitutions (Extended data Figs 5b, c and 6b, c), suggesting that the B.1.351.3 mutations were selected by antibody pressure.

## A mutant SARS-CoV-2 spike with high-level plasma resistance

Based on the above findings, we attempted to generate a mutant SARS-CoV-2 spike protein that was minimally divergent from SARS-CoV-2 Wuhan-hu-1, yet resistant to neutralization by polyclonal convalescent and vaccine recipient plasma. Successful derivation of such a spike protein would identify a complete list of neutralization epitopes recognized by polyclonal antibodies. We chose 20 naturally occurring mutations, including 8 NTD and 8 RBD changes (Fig. 3d) that either (i) arose in our plasma selection experiments (Fig. 2b), (ii) occur in VOC with reduced neutralization sensitivity (Extended data Fig. 5a, b) or (iii) arose in our previous studies where human monoclonal antibody resistance was selected^4,14,24^. Naturally occurring deletion mutations in the NTD (Extended data Fig. 5a), as well as multiple substitutions confering resistance to class 1, 2,and 3 RBD-binding antibodies^4,14,24^ were included. An rVSV/SARS-CoV-2 derivative encoding the resulting spike sequence, termed PMS20 (Fig. 3d) was replication competent but attenuated compared to rVSV/SARS-CoV-2_2E1_, suggesting that the 20 mutations confer a fitness cost (Fig 3e). Nevertheless, HIV-1 pseudotypes bearing PMS20, were similarly infectious to those bearing the parental spike protein (Extended Data Fig. 1) and were highly resistant to neutralization. Indeed, 17/21 random convalescent and 8/15 mRNA vaccinee plasmas gave undetectable neutralization of PMS20 pseudotypes (<1:50, Fig 3f). Among the high-titer convalescent RU27 plasmas 23/27 had residual neutralizing activity against PMS20 that was reduced by a median of 32-fold compared to the parental pseudotype (range 2.8 – 114 fold, Extended Data Fig. 6a). We conclude that the 20 mutations in the PMS20 spike protein are sufficient for evasion of the majority of the antibodies in the plasma of individuals who have been infected by or vaccinated against SARS-CoV-2.

## Polyclonal neutralization breadth in vaccinated convalescents

However, a panel of plasmas from individuals who had been both infected and subsequently received mRNA vaccines^25^ retained neutralizing activity against HIV-1 pseudotypes bearing the PMS20 spike (Fig 4a, b). Indeed, the PMS20 mutations that reduced Ran21 and Vac14 plasma NT_50_ by a median of 50-fold (range 5.9 to 225 fold) and 81-fold (range 8.4 to 229-fold), respectively caused a median NT_50_ reduction of only 18.6-fold (range 3.9 to 100-fold) for the vaccinated convalescent (VC1-15) plasma panel (Fig 4b). Analysis of chimeric SARS-CoV-2/PMS20 spike proteins in which the respective RBDs were exchanged [PMS20(2-RBD) and SARS-CoV-2(PMS-RBD)] indicated that the relative resistance of the PMS20 to both Ran and VC plasmas was conferred by multiple spike determinants and that the neutralization breadth in the VC plasmas was due to antibodies directed at both RBD and non-RBD determinants (Fig. 4a). In addition to the previously reported potent neutralizing activity of VC plasmas against the B.1.1.7, B.1.525, P.1 and B.1.351.3 VOCs^25^, the VC plasmas also potently neutralized B.1.617.2 (delta), as well as a recently described variant (A.VOI.V2)^26^ that has 11 substitutions and 3 deletions in spike, including an extensively mutated NTD, and is predicted to be resistant to both class 2 and class 3 RBD-binding neutralizing antibodies (Extended Data Fig. 7).

Plasma from the vaccinated/convalescent group also had substantial neutralizing activity against heterologous sarbecovirus HIV-1 pseudotypes, including those that were poorly neutralized by Ran21, Vac14 and RU27 plasma panels and whose RBD and/or NTD sequences are extensively divergent from SARS-CoV-2 (Fig 4c, d). The median NT_50_ values for the VC plasmas against sarbecovirus pseudotypes were 5330 (range 2369-7222) for bCoV RaTG13; 3617 (range 1780-6968) for pCoV-GX; 2605 (range 1386-3181) for bCoV-WIV16 and 1208 (range 621-2705) SARS-CoV (Fig. 4d, Extended Data Fig. 7). Notably, the neutralizing activity of the VC plasmas against the highly divergent sarbecoviruses bCoV-WIV16 and SARS-CoV (Fig. 4c) was similar to that found in the random convalescent plasmas against SARS-CoV-2 Wuhan-hu-1 (Extended Data Fig. 7), Thus, the neutralization potency and breadth of polyclonal plasma following mRNA vaccination of previously SARS-CoV-2 infected individuals appears greater than previously appreciated.

## Discussion

The sensitivity of chimeric and polymutant spike proteins and a requirement for numerous mutations in the acquisition of resistance to plasma neutralization, indicates abundant neutralizing antibody targets on the SARS-CoV-2 spike protein. Thus, there is a high genetic barrier to complete escape from the polyclonal neutralizing antibodies generated by randomly selected convalescents and mRNA vaccine recipients. Additionally, our recent analyses suggests that affinity maturation, over months, of SARS-CoV-2 neutralizing antibodies in convalescents confers additional flexibility and affinity^24,25,27^ that may not be afforded by standard mRNA vaccine regimens^28^. Indeed, individual affinity-matured antibodies can impose a requirement for multiple viral substitutions for antibody escape and enable substantial activity against VOC. Some human monoclonal antibodies also have activity against divergent sarbecoviruses^24,29^. Thus, affinity maturation, the availability of numerous epitope targets and the generation of high levels of circulating antibodies may explain why polyclonal plasma from individuals who have been both infected and subsequently vaccinated could effectively neutralize the otherwise highly neutralization resistant PMS20 spike, as well as sarbecoviruses whose NTD and/or RBD domains are divergent from SARS-CoV-2. It remains to be seen whether similar neutralization potency and breadth can be achieved using appropriately timed boosting with existing SARS-CoV-2 vaccines. If so, existing immunogens may be sufficient to provide robust protection against SARS-CoV-2 variants that may arise in future, and a degree of protection against potential future sarbecovirus threats. Conversely, PMS proteins encoding numerous neutralization escape mutations may represent useful immunogens to broaden the polyclonal antibody response elicited by first generation SARS-CoV-2 vaccines.

## Data availability

The datasets generated during and/or analyzed during the current study are available in the accompanying source data files and from the corresponding author on reasonable request

## Methods

### SARS-CoV-2 pseudotyped reporter virus

Plasmids pSARS-CoV-2-SΔ19 and pSARS-CoV-SΔ19 expressing C-terminally truncated SARS-CoV-2 (NC_045512) and SARS-CoV spike proteins have been described previously^19^ and were used to construct the SARS-CoV-2(1-RBD) and SARS-CoV(2-RBD) expression plasmids in which RBD-encoding sequences were reciprocally exchanged. A panel of plasmids expressing spike proteins from SARS-CoV-2 VOC and VOI were constructed in the context of pSARS-CoV-2-SΔ19 (R683G) ^19^. Substitutions were introduced using synthetic gene fragments (IDT) or overlap extension PCR mediated mutagenesis and Gibson assembly. All VOC/VOI and polymutant spike proteins also included the R683G substitution, which disrupts the furin cleavage site and generates higher titer virus stocks without significant effects on pseudotyped virus neutralization sensitivity (Extended Data Fig. 1c, d). The potencies with which the plasma neutralized members of the mutant pseudotype panel were compared with potencies against a “wildtype” SARS-CoV-2 spike sequence, carrying R683G where appropriate. Plasmids expressing the spike proteins found in the horseshoe bat (*Rinolophus affinis*) coronavirus bCoV-RaTG13 ^21^ as well as the pangolin (*Manis javanica*) coronaviruses from Guandong, China (pCoV-GD) and Guanxi, China (pCoV-GX) ^22,23^ were similarly constructed. Spike sequences were codon-modified to maximize homology with the human codon-usage optimized of the pSARS-CoV-2 expressing plasmid VG40589-UT (Sinobiological). The 19aa truncated CDS of bCoV-RaTG13 (QHR63300), pCoV-GD (CoV_EPI_ISL_410721), and pCoV-GX (CoV_EPI_ISL_410542) were synthesized by GeneART and subcloned into pCR3.1 using NheI and XbaI and Gibson assembly, and referred to as pCR3.1-bCoV-RaTG13-SΔ19, pCR3.1pCoV-GD-SΔ19 and pCR3.1-pCoV-GX-SΔ19, respectively. Pseudotyped HIV-1 particles were generated as previously described^19^. Specifically, virus stocks were harvested 48 hours after transfection of 293T cells with pHIV-1 GagPol and pCCNano/LucGFP (Fig.1) or pNL4-3ΔEnv-nanoluc (all other Figs) along with a spike expression plasmid, filtered and stored at −80°C.

### SARS-CoV2-2/sarbecovirus pseudotype neutralization assays

Plasmas were five-fold serially diluted and then incubated with pseudotyped HIV-1 reporter virus for 1 h at 37 °C. The antibody/pseudotype virus mixture was then added to HT1080/ACE2.cl14 cells. After 48 h, cells were washed with PBS, lysed with Luciferase Cell Culture Lysis reagent (Promega) and Nanoluc Luciferase activity in lysates was measured using the Nano-Glo Luciferase Assay System (Promega) and a Glomax Navigator luminometer (Promega). The relative luminescence units were normalized to those derived from cells infected with the pseudotyped virus in the absence of plasma. The half-maximal neutralizing titer (NT50) was determined using four-parameter nonlinear regression (least squares regression method without weighting) (GraphPad Prism).

### Plasma samples

Plasma samples were from individuals who were infected with SARS-CoV2 a mean of 1.3 months prior to plasma donation^2^ or from individuals who had received mRNA vaccines at various times prior to plasma donation ^4^. A set of twenty-seven plasmas samples from SARS-CoV-2 infected individuals with high neutralizing activity who had not been vaccinated^2^, termed the “RU27” plasma panel were used in VSV-SARS-CoV-2 selection procedures, while this panel plus a second set of 21 randomly selected plasmas (selected at random with blinding to neutralization titer or any demographic characteristic) from the same convalescent cohort formed the “Ran21” plasma panel ^2^. A set of 14 plasmas donated by individuals who had received a Pfizer/BionTech mRNA vaccine formed the “Vac14” plasma panel ^4^. Finally a set of 15 plasmas from convalescent individuals who had received a Pfizer/BionTech mRNA vaccine between 6 and 12 months after infection ^25^ formed the “VC15” plasma panel. The study visits and blood draws were reviewed and approved by the Institutional Review Board of the Rockefeller University (IRB no. DRO-1006, ‘Peripheral Blood of Coronavirus Survivors to Identify Virus-Neutralizing Antibodies’).

### Selection of antibody resistant rVSV/SARS-CoV-2 variants

To select plasma-resistant spike variants, rVSV/SARS-CoV-2/GFP1D7 and rVSV/SARS-CoV-2/GFP_2E1_ were passaged to generate diversity, and populations containing 10^6^ PFU were incubated with plasma (diluted 1:50 to 1:400) for 1h at 37°C before inoculation of 2×10^5^ 293T/ACE2cl.22 cells in 6-well plates. The following day the medium was replaced with fresh medium containing the same concentrations of plasma. Supernatant from the wells containing the highest concentrations of plasma antibodies that showed evidence of rVSV/SARS-CoV-2/GFP replication (large numbers of GFP positive cells or GFP positive foci) was harvested 24h later. Thereafter, aliquots (100 μl) of the cleared supernatant from the first passage (p1) were incubated with the same or increased concentration of plasma and then used to infect 2×10^5^ 293T/ACE2cl.22 cells in 6-well plates, as before (p2). In situations where small, but expanding GFP-positive foci were observed, the medium was refreshed at 48h with fresh medium containing no plasma and the virus harvested 24h later. We repeated this process for up to 6 passages or until reduced neutralization potency for the plasma was obvious, as indicated by visual detection of increasing numbers of GFP positive cells during passage.

To isolate individual mutant viruses by limiting dilution, the selected rVSV/SARS-CoV-2/GFP_1D7_ and rVSV/SARS-CoV-2/GFP_2E1_ populations were serially diluted in the absence of plasma and aliquots of each dilution added to individual wells of 96-well plates containing 1×10^4^ 293T/ACE2cl.22 cells. Individual viral variants were identified by observing single GFP-positive plaques in individual wells at limiting dilutions. The plaque-purified viruses were expanded, RNA was extracted, and spike sequences determined.

### rVSV/SARS-CoV-2 Neutralization assays

Plasma samples were five-fold serially diluted and then incubated with rVSV/SARS-CoV-2/GFP_1D7_ and rVSV/SARS-CoV-2/GFP_2E1_, or plaque purified selected variants thereof, for 1 h at 37 °C. The antibody/recombinant virus mixture was then added to 293T/ACE2.cl22 cells. After 16h, cells were harvested, and infected cells were quantified by flow cytometry. The percentage of infected cells was normalized to that derived from cells infected with rVSV/SARS-CoV-2 in the absence of plasma. The half-maximal neutralizing titer for each plasma (NT_50_) was determined using four-parameter nonlinear regression (least squares regression method without weighting) (GraphPad Prism).

### Sequence analyses

To identify putative antibody resistance mutations, RNA was isolated from aliquots of supernatant containing selected viral populations or individual plaque purified variants using NucleoSpin 96 Virus Core Kit (Macherey-Nagel). The purified RNA was subjected to reverse transcription using random hexamer primers and SuperScript VILO cDNA Synthesis Kit (Thermo Fisher Scientific). The cDNA was amplified using KOD Xtreme Hot Start DNA Polymerase (Millipore Sigma). Specifically, a fragment including the coding region of the extracellular domain of spike was amplified using primers targeting the intergenic region between VSV-M and spike, and the spike intracellular domain. The PCR products were purified and sequenced either using Sanger-sequencing or Illumina sequencing as previously described ^30^. For Illumina sequencing, 1 μl of diluted DNA was used with 0.25 μl Nextera TDE1 Tagment DNA enzyme (catalog no. 15027865), and 1.25 μl TD Tagment DNA buffer (catalog no. 15027866; Illumina). Then, the DNA was ligated to i5/i7 barcoded primers using the Illumina Nextera XT Index Kit v2 and KAPA HiFi HotStart ReadyMix (2X; KAPA Biosystems). Next the DNA was purified using AmPure Beads XP (Agencourt), pooled, sequenced (paired end) using Illumina MiSeq Nano 300 V2 cycle kits (Illumina) at a concentration of 12pM.

For analysis of the Illumina sequencing data, adapter sequences were removed from the raw reads and low-quality reads (Phred quality score <20) using BBDuk. Filtered reads were mapped to the codon-optimized SARS-CoV-2 S sequence in rVSV/SARS-CoV-2/GFP and mutations were annotated using using Geneious Prime (Version 2020.1.2), using a P-value cutoff of 10^−6^. RBD-specific variant frequencies, P-values, and read depth were compiled using Python running pandas (1.0.5), numpy (1.18.5), and matplotlib (3.2.2). The parental rVSV/SARS-CoV-2/GFP 2E1 and 1D7 sequences each conatain two adaptive mutations(1D7, F157S and R685M for 1D7; D215G and R683G for 2E1) but each was considered “WT” for the purposes of the plasma selection experiments and were subtracted from the analyses of the sequences.

The frequency of amino acid substitutions during rVSV/SARS-CoV-2 passage in plasmas was compared with the frequency of global occurrences of changes at each residue on 5/11/21 (Los Alamos, COVID-19 Viral Genome Analysis Pipeline, https://cov.lanl.gov/content/index)^31^. For comparison of SARS-CoV-2 with sarbecoviruses, amino acid sequences were aligned with Clustal Omega. Using a python script clone of Simplot (https://jonathanrd.com/20-05-02-writing-a-simplot-clone-in-python/), the percent identity relative to SARS-CoV2 was calculated within a rolling window of 100 amino acids, stepping a single residue at a time.

For three-dimensional sliding window analysis of changes in the spike amino acid sequence observed globally and in vitro, the frequency of global occurrences of changes at each residue (Los Alamos, COVID-19 Viral Genome Analysis Pipeline, https://cov.lanl.gov/content/index) ^31^was divided by the average frequency of change at any reside and projected in the SARS-CoV-2 spike structure PDB 6VXX ^32^ as relative change frequency using BioStructMap ^33,34^. Alternatively, the averaged frequency of substitutions observed after passaging rVSV/SARS-CoV-2 with plasma was divided by the mean substitution frequency and applied as a 3D sliding window over the spike structure. The average frequency of substitutions in a 15 Å radius is represented using a color spectrum.

#### Acknowledgements

This work was supported by NIH grant R37AI64003 and R01AI501111 (P.D.B).; R01AI78788 (TH); P01-AI138398-S1 (M.C.N.) and 2U19AI111825 (M.C.N.). C.G. was supported by the Robert S. Wennett Post-Doctoral Fellowship, in part by the National Center for Advancing Translational Sciences (National Institutes of Health Clinical and Translational Science Award programme, grant UL1 TR001866), and by the Shapiro–Silverberg Fund for the Advancement of Translational Research. P.D.B. and M.C.N. are Howard Hughes Medical Institute Investigators.

#### Author contributions

P.D.B., T.H., M.C.N., FS., and Y.W. conceived, designed and analyzed the experiments. F.S., Y.W., constructed and performed rVSV/SARS-CoV-2 selection and neutralization experiments. F.S., Y.W., M.R., J.D.S, and E.B. performed pseudotype neutralization experiments. F.S., T.H. and F.Z. constructed expression plasmids. A.C. performed NGS. D.P. performed bioinformatic analysis. M.C, C.G. and D. J. S-B executed clinical protocols and recruited participants and processed samples. P.D.B., T.H., FS., and Y.W. wrote the manuscript with input from all co-authors.

---

**Extended Data Fig. 1.**
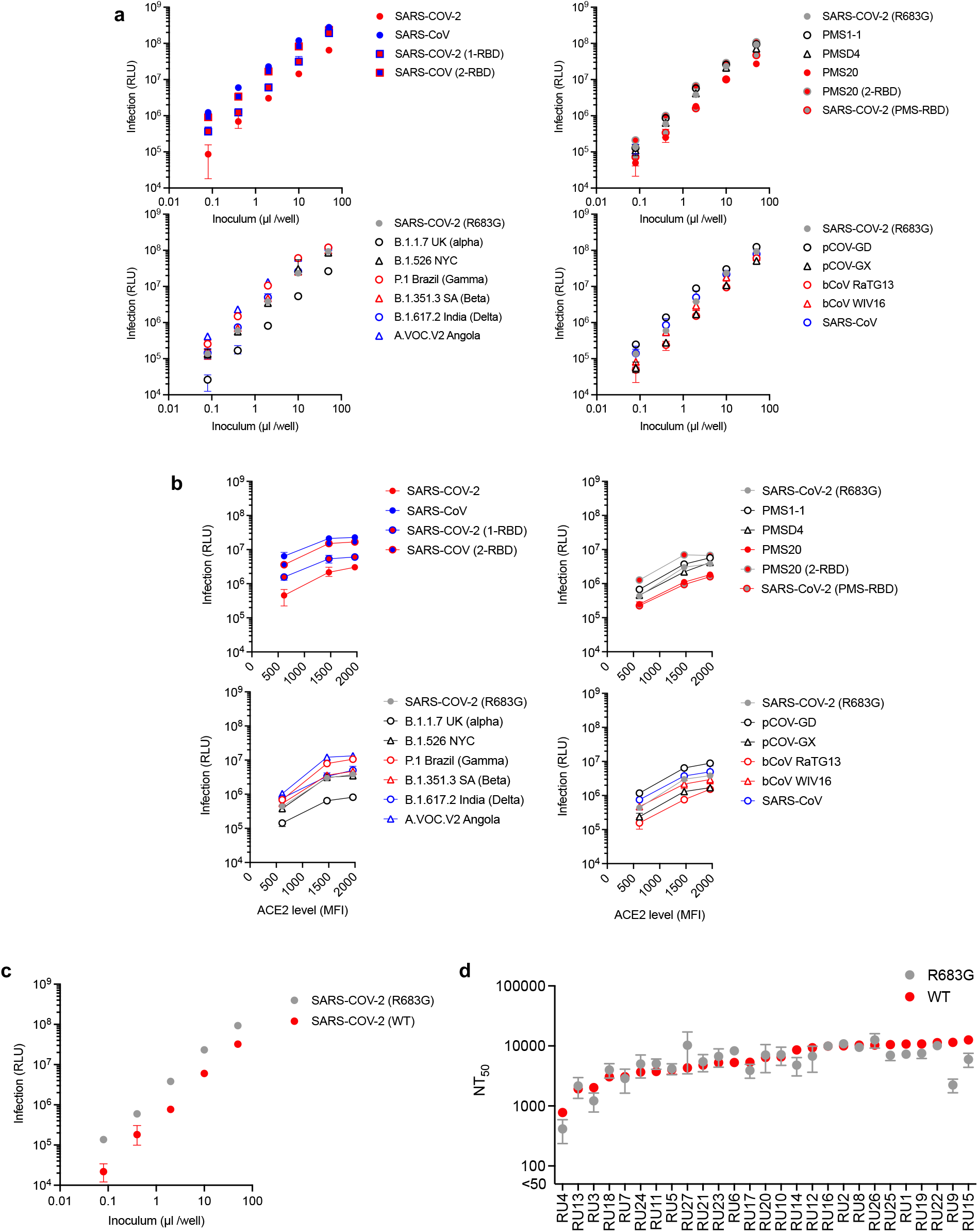
Characterization of HIV-1 pseudotypes bearing the chimeric, mutant, and variant SARS-CoV-2 and sarbecovirus spike proteins. (**a**) Titration of pseudotyped viruses on 293T/ACE2cl.22 cells. Chimeric spike pseudotyped viruses in the upper left panel were built using the unaltered SARS-CoV-2 and SARS-CoV spike proteins and a 3-plasmid HIV-1 pseudotyping system (see Methods). The other panels depict titration of pseudotypes derived using a furin cleavage site mutant SARS-CoV-2 spike protein (R683G) and a 2-plasmid HIV-1 pseudotyping system (see Methods). (**b**) The same pseudotyped viruses used in (**a**) were used to infect 3 different 293T/ACE2 clonal cell lines each expressing a different level of ACE2 (MFI = mean fluorescence intensity). (**c**) Titration of pseudotypes derived bearing an unaltered SARS-CoV-2 spike protein and a furin cleavage site mutant SARS-CoV-2 spike protein (R683G) generated using a 2-plasmid HIV-1 pseudotype system (see Methods). (**d**) Comparative neutralization potency (NT_50_ values) of high titer convalescent (RU27) plasmas against HIV-1 pseudotypes bearing R683G mutant (grey symbols) and unaltered (red symbols) SARS-CoV-2 spike proteins.

**Extended Data Fig. 2.**
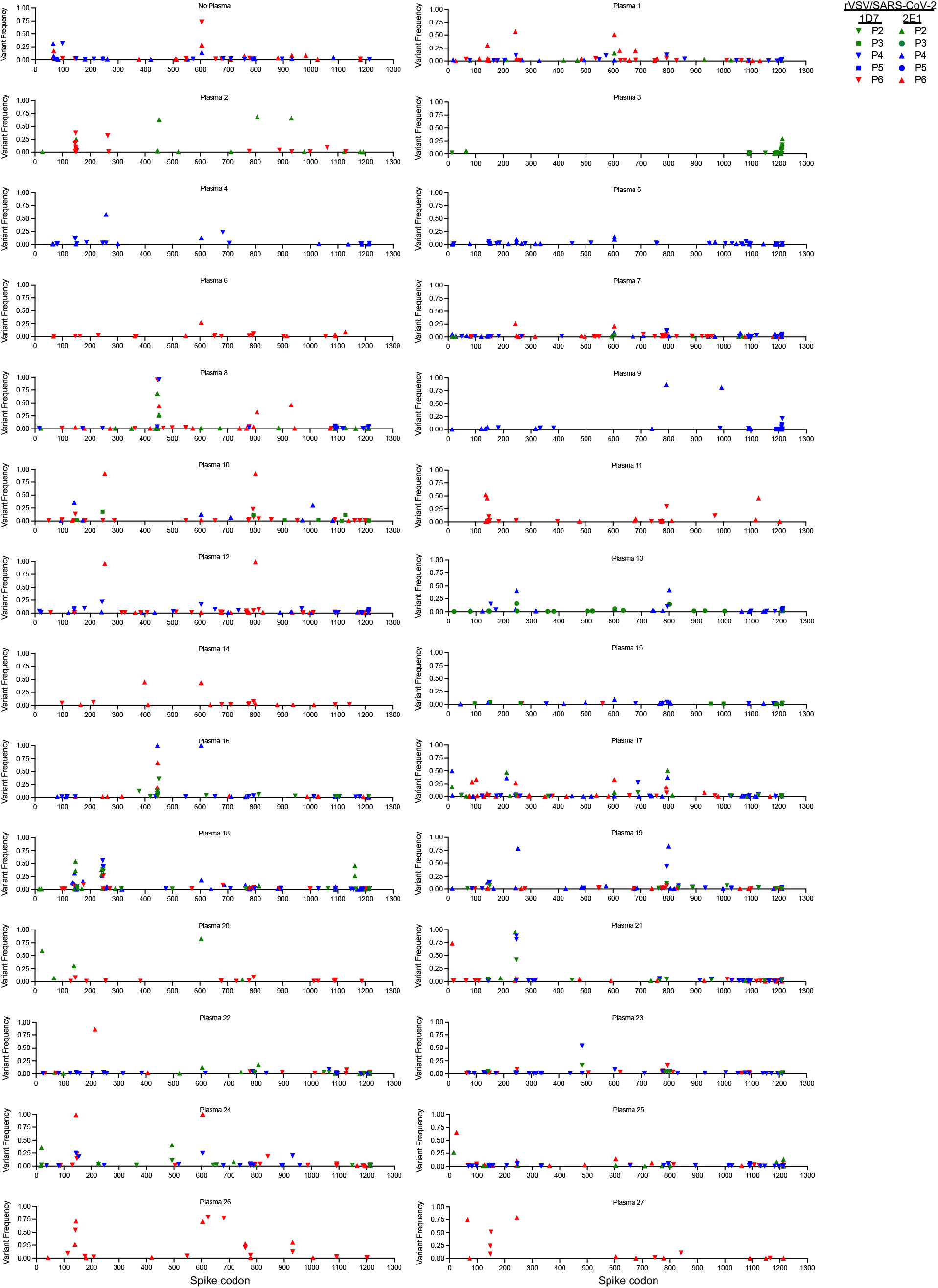
Selection pressure on SARS-CoV-2 spike exerted by convalescent plasma. Frequencies of amino acid substitutions at each codon of the SARS-CoV-2 spike protein following the indicated number of passages (P2-P6) of two independent rVSV-SARS-CoV-2 populations (1D7 and 2E1), in each of the RU27 plasmas, determined by NGS sequencing.

**Extended Data Fig. 3.**
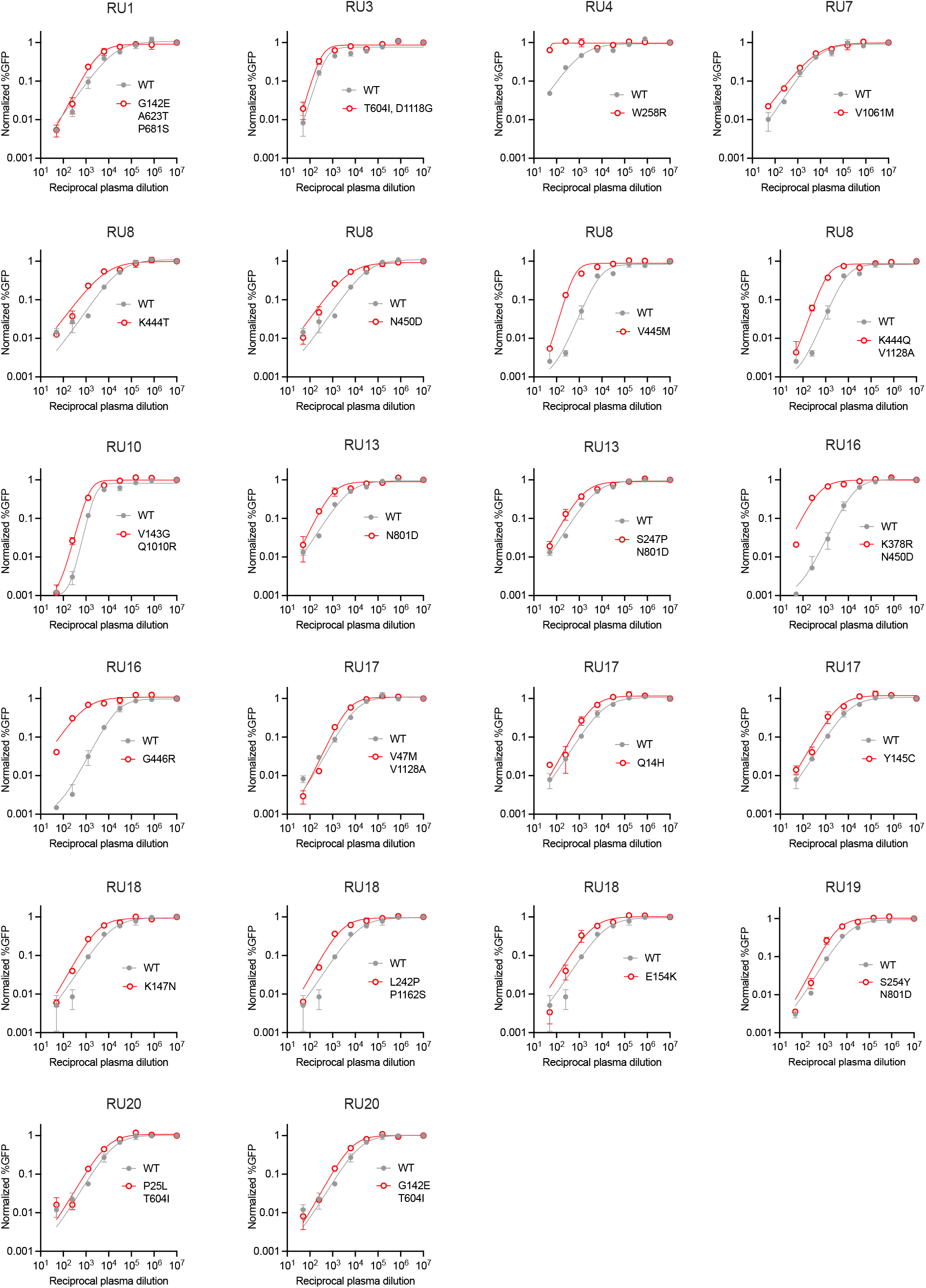
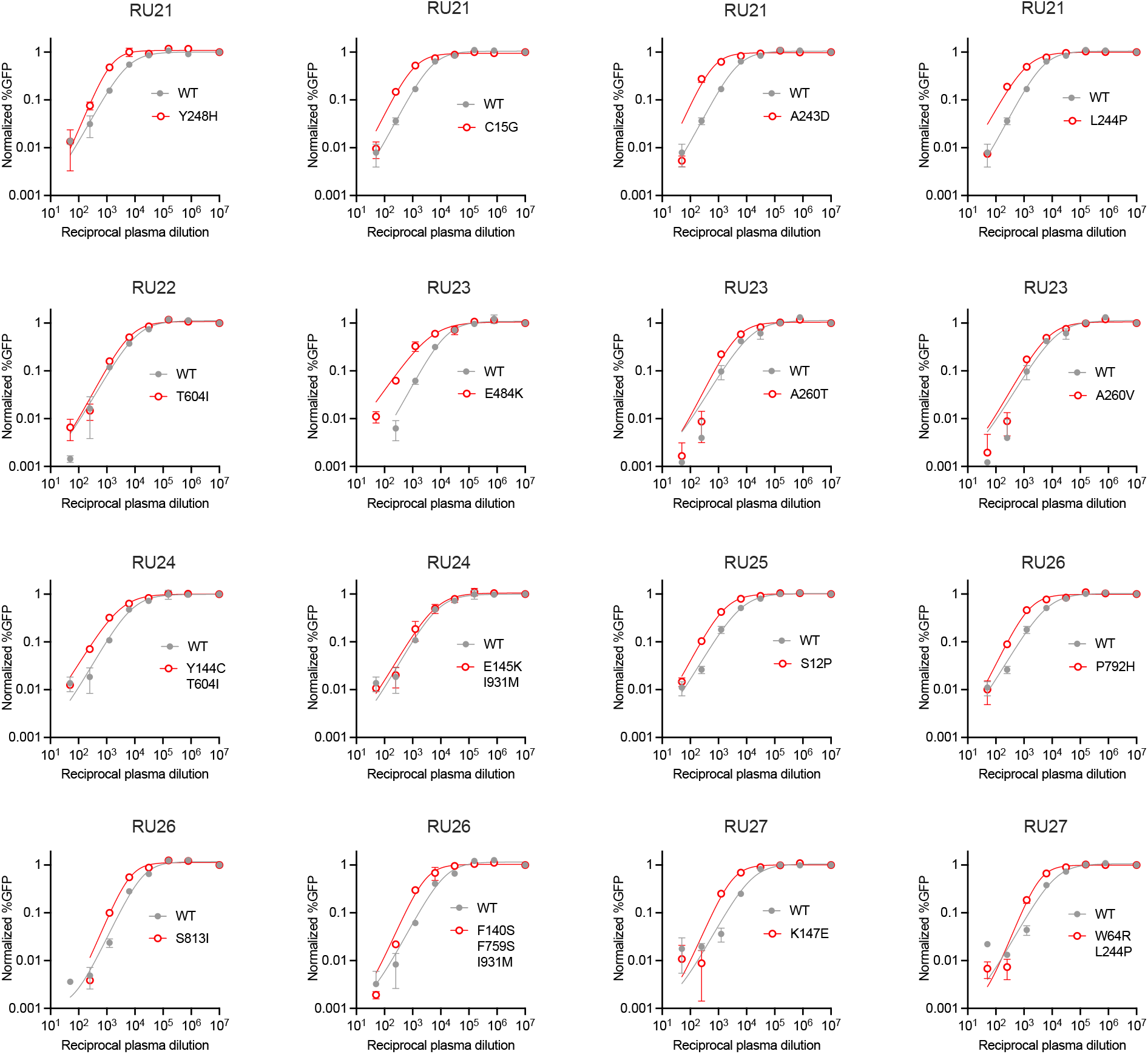
Neutralization sensitivity of plasma-selected rVSV/SARS-CoV-2 mutants. Infection, relative to non-neutralized controls, by plaque purified rVSV/SARS-CoV-2 isolates in the presence of the indicated dilutions of the indicated plasmas from the RU27 panel. The same plasmas that were used to select the indicated mutants were used to determine neutralization potency against the respective plaque purified mutants (red) and parental (WT, grey) rVSV/SARS-CoV-2 1D7 or 2E1 viruses. Median and range of two technical replicates is plotted representative of two independent experiments.

**Extended data Fig. 4.**
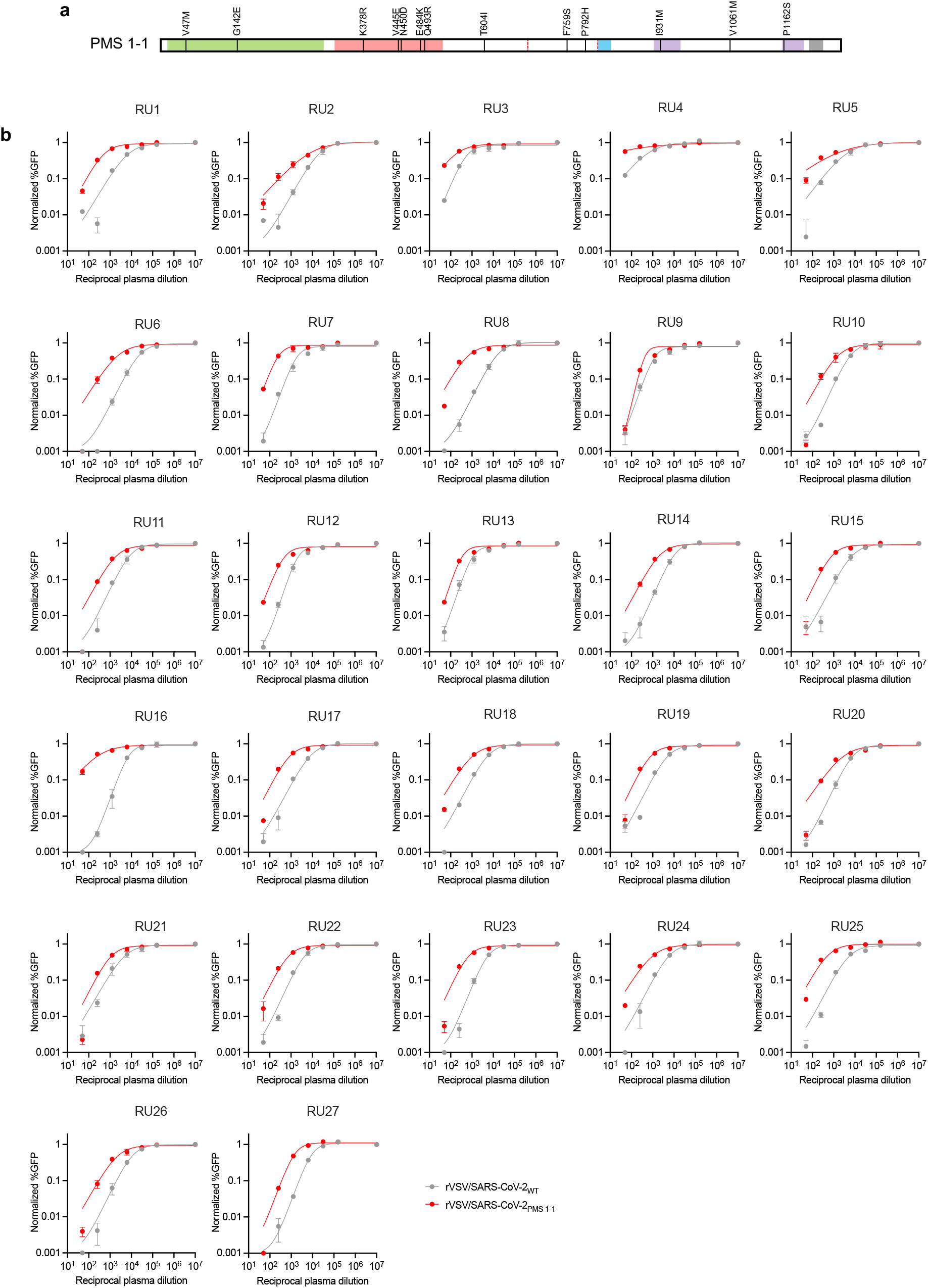
Neutralization sensitivity of rVSV/SARS-CoV-2 encoding the PMS1-1 spike. (**a**) Design of the PMS1-1 polymutant spike protein with 13 plasma-selected spike mutations aggregated in a single spike. (**b**) Infection, relative to non-neutralized controls, by rVSV/SARS-CoV2_PMS1-1_ (red) and rVSV/SARS-CoV2_2E1_ (grey) in the presence on the indicated dilutions of the plasmas from the RU27 panel. Median and range of two technical replicates is plotted, representative of two independent experiments.

**Extended Data Fig 5.**
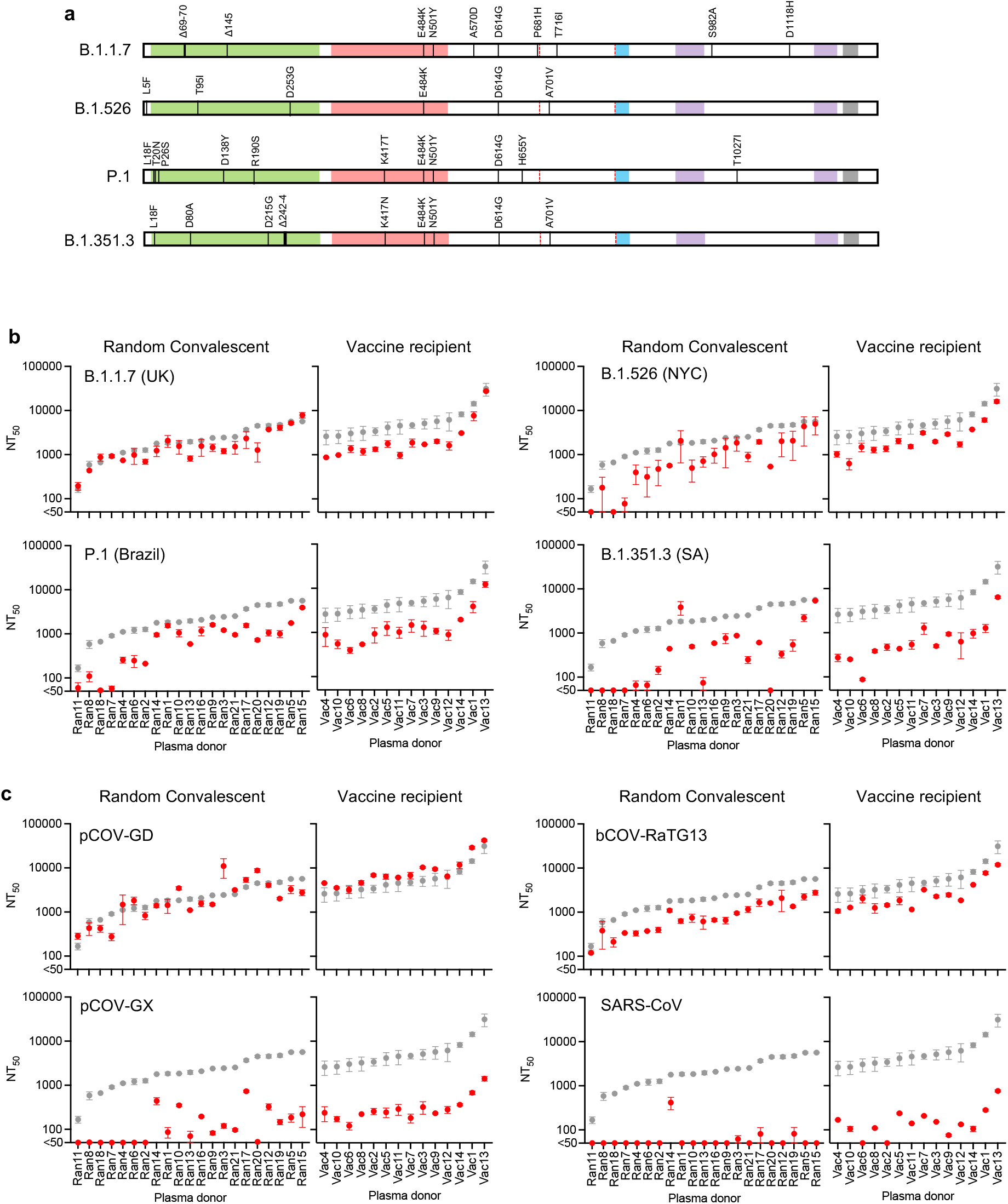
Neutralization potency of random convalescent and vaccine recipient plasmas against VOC/VOI, and sarbecovirus HIV-1 pseudotypes. (**a**) Schematic representation of substitutions in naturally occurring VOC/VOI SARS-CoV-2 spike proteins. (**b**,**c**) Comparative neutralization potency (NT_50_ values) of random convalescent (Ran1-21) and vaccine recipient (Vac1-14) plasmas plasma against WT (grey symbols) and indicated SARS-CoV-2 variant (**b**) or sarbecovirus (**c**) (red symbol) HIV-1 pseudotypes. Median and range of two independent experiments, each with two technical replicates, is plotted.

**Extended Data Fig 6.**
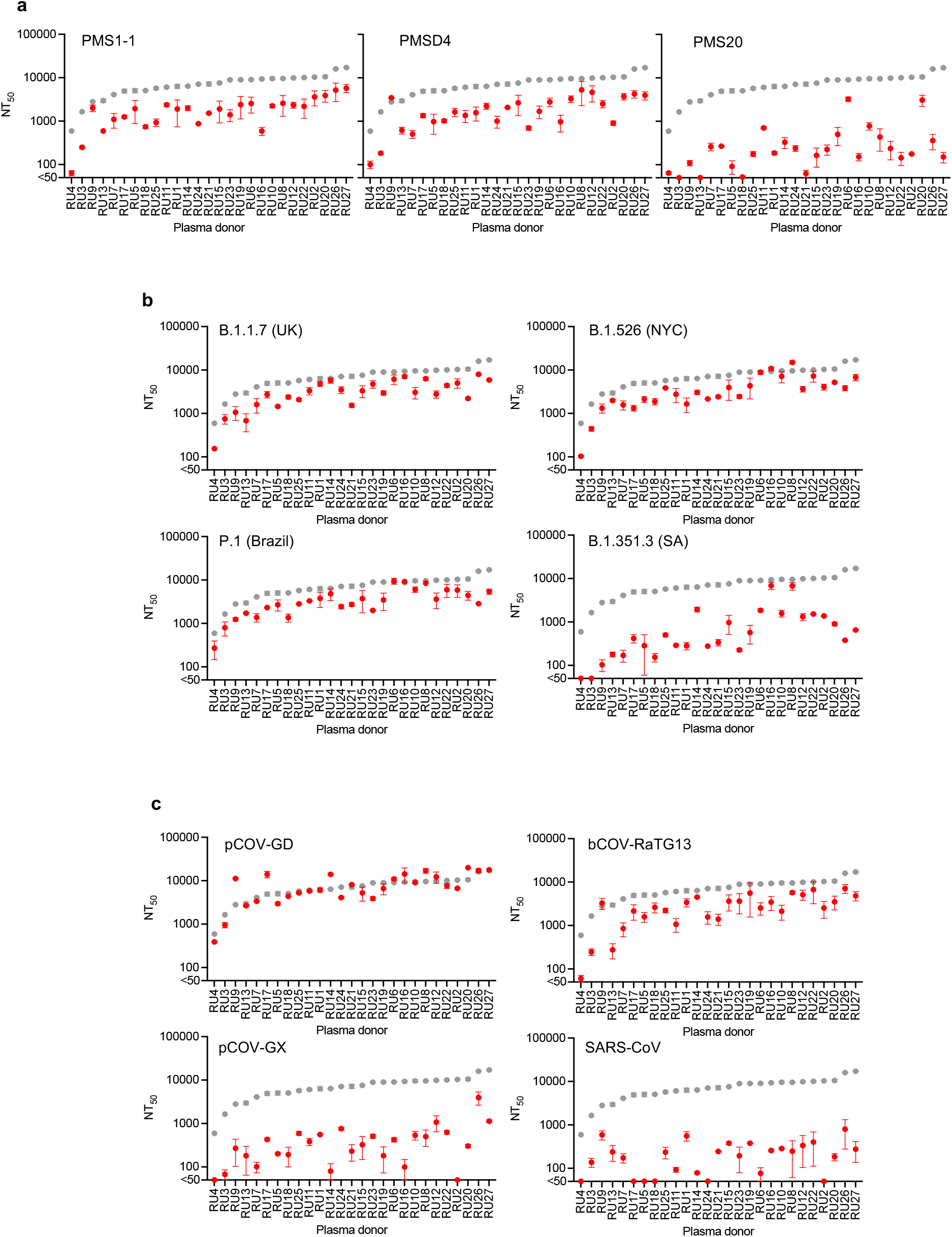
Neutralization potency of high titer convalescent plasma against PMS, VOC/VOI, and sarbecovirus HIV-1 pseudotypes. (**a**-**c**) Comparative neutralization potency (NT_50_ values) of high titer convalescent (RU27) plasma against WT (grey symbols) and indicated PMS (**a**), SARS-CoV-2 variant (**b**) or sarbecovirus (**c**) (red symbol) HIV-1 pseudotypes. Median and range of two independent experiments, each with two technical replicates, is plotted.

**Extended Data Fig 7.**
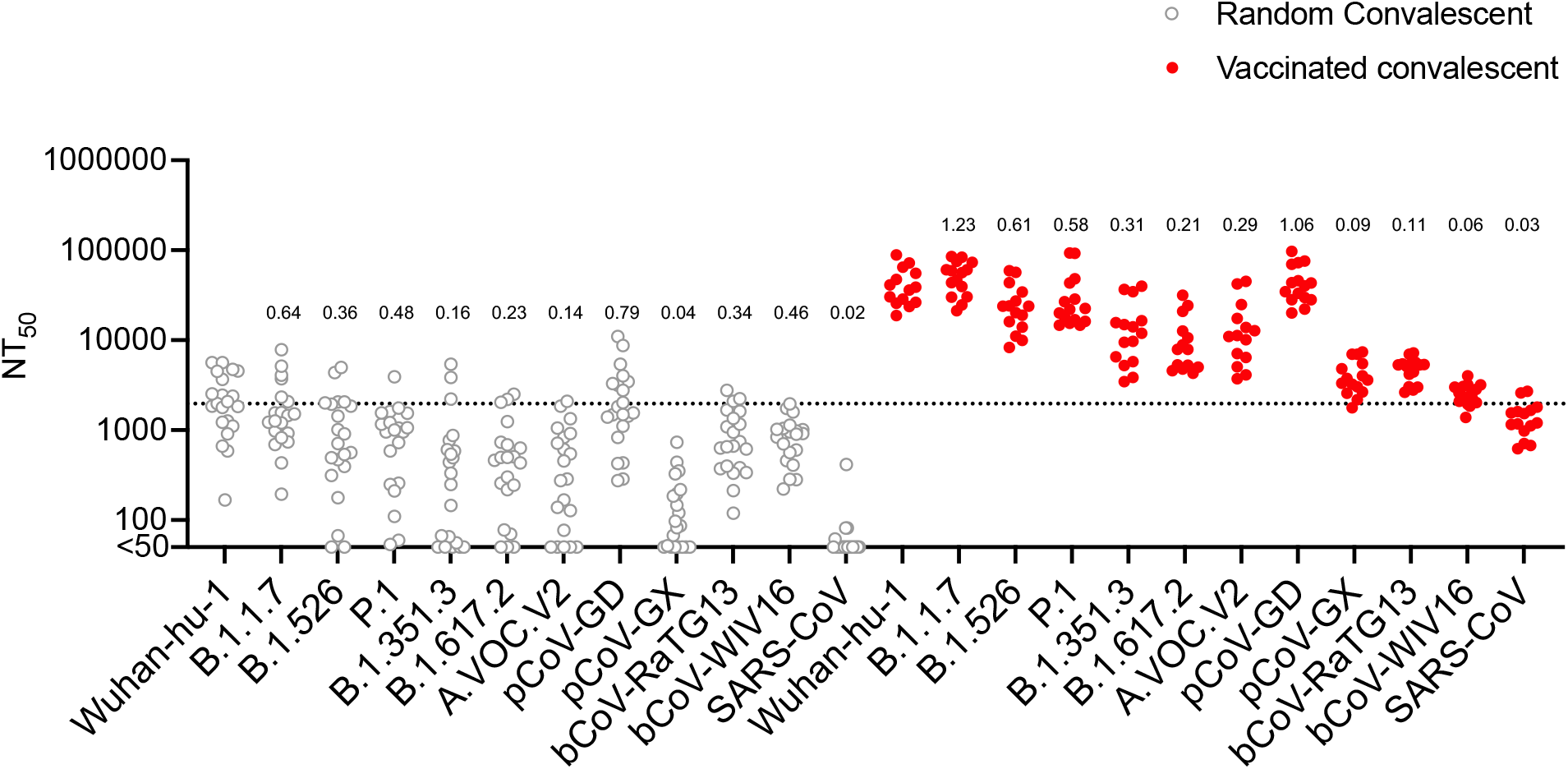
Neutralization potency of plasma from vaccinated convalescents against VOC/VOI and sarbecovirus HIV-1 pseudotypes. Neutralization potency (NT_50_ values) of random convalescent plasmas (grey symbols) against or vaccinated convalescents’ plasma (red symbols) against SARS-CoV-2 prototype or variant or sarbecovirus HIV-1pseudotypes. Median of two independent experiments, each with two technical replicates, is plotted. Numbers above each scatterplot indicate the median NT_50_ relative to the median NT_50_ for Wuhan-Hu-1 SARS-CoV-2.

**Table S1.** Substitutions enriched in rVSV/SARS-CoV-2 following selection in neutralizing plasma.

| Plasma | Parental virus | Dilution | Passage | Substitutions |
| --- | --- | --- | --- | --- |
| RU1 | rVSV/SARS-CoV-2<sub>1D7</sub> | 100 | 4 | I794T (11.7%), R246G (10.9%), T573I (10.3%) |
|  | rVSV/SARS-CoV-2<sub>1D7</sub> | 100 | 6 | I794T (5.7%), N536H (5.1%) |
|  | rVSV/SARS-CoV-2<sub>2E1</sub> | 100 | 2 | T604I (15.4%) |
|  | rVSV/SARS-CoV-2<sub>2E1</sub> | 200 | 6 | L244P (57.2%), T604I (50.8%), G142E (30.7%), A623T (20.3%), P681S (20%), F759S (5.7%), Y248H (5.2%) |
| RU2 | rVSV/SARS-CoV-2<sub>1D7</sub> | 200 | 6 | N148K (37.3%), A264D (32.2%), N149T (19.5%), H146R (16.8%), N148S (9.6%), V1061A (8.7%), K150N (5.9%) |
|  | rVSV/SARS-CoV-2<sub>2E1</sub> | 200 | 6 | P807H (68.5%), I931M (65.9%), N450D (63.2%), K150N (25.6%) |
| RU3 | rVSV/SARS-CoV-2<sub>1D7</sub> | 50 | 2 | P1213S (17.6%), W1212R (10.2%), K1211R (8.8%), I1210N (6%) |
|  | rVSV/SARS-CoV-2<sub>2E1</sub> | 50 | 2 | P1213S (29.5%), W1212R (19%), K1211R (16.2%), I1210N (12%), W64R (6%) |
| RU4 | rVSV/SARS-CoV-2<sub>1D7</sub> | 50 | 4 | R682G (23.7%), Y145D (12.5%), K147R (11.8%) |
|  | rVSV/SARS-CoV-2<sub>2E1</sub> | 50 | 4 | W258R (58.3%), T604I (12.8%) |
| RU5 | rVSV/SARS-CoV-2<sub>1D7</sub> | 50 | 4 | K147E (6.4%), I1081V (5.4%) |
|  | rVSV/SARS-CoV-2<sub>2E1</sub> | 50 | 4 | T604I (10%), Y248D (5.4%) |
|  | rVSV/SARS-CoV-2<sub>2E1</sub> | 50 | 4 | T604I (14.9%), Y248D (10.5%) |
| RU6 | rVSV/SARS-CoV-2<sub>1D7</sub> | 400 | 6 | I794T (6%) |
|  | rVSV/SARS-CoV-2<sub>2E1</sub> | 400 | 6 | T604I (27.2%), V1128A (9.2%) |
| RU7 | rVSV/SARS-CoV-2<sub>1D7</sub> | 50 | 4 | I794T (11.6%), P1213S (6.1%) |
|  | rVSV/SARS-CoV-2<sub>1D7</sub> | 200 | 4 | I794T (13%) |
|  | rVSV/SARS-CoV-2<sub>1D7</sub> | 100 | 6 | N709D (5.6%), I794T (5.3%) |
|  | rVSV/SARS-CoV-2<sub>2E1</sub> | 100 | 3 | T604I (6.5%) |
|  | rVSV/SARS-CoV-2<sub>2E1</sub> | 100 | 4 | T604I (9.3%), V1061M (8.5%), E1188D (6.4%), C15Y (6%) |
|  | rVSV/SARS-CoV-2<sub>2E1</sub> | 100 | 6 | L244P (26.5%), T604I (21.4%), I794T (6.1%), R1091S (5.1%) |
| RU8 | rVSV/SARS-CoV-2<sub>1D7</sub> | 50 | 4 | N450D (95.2%), R1091S (5.5%) |
|  | rVSV/SARS-CoV-2<sub>1D7</sub> | 200 | 6 | V445M (94.8%) |
|  | rVSV/SARS-CoV-2<sub>2E1</sub> | 50 | 4 | K444T (67.9%), N450D (26.3%) |
|  | rVSV/SARS-CoV-2<sub>2E1</sub> | 100 | 4 | K444T (68.4%), N450D (27.8%) |
|  | rVSV/SARS-CoV-2<sub>2E1</sub> | 200 | 6 | I931M (46.1%), N450D (44.1%), P807H (32.6%) |
| RU9 | rVSV/SARS-CoV-2<sub>1D7</sub> | 50 | 4 | P1213S (21%), W1212R (9.7%), K1211R (8.5%), I1210N (5.1%) |
|  | rVSV/SARS-CoV-2<sub>2E1</sub> | 100 | 4 | P792H (86.5%), Q992H (81%), P1213S (6%) |
| RU10 | rVSV/SARS-CoV-2<sub>1D7</sub> | 100 | 3 | R246G (18%), I794T (11.7%), V1128M (11.6%) |
|  | rVSV/SARS-CoV-2<sub>1D7</sub> | 200 | 6 | P792H (22.6%), K147R (13.3%), I794T (7.3%) |
|  | rVSV/SARS-CoV-2<sub>2E1</sub> | 50 | 4 | V143G (35.8%), Q1010R (30.5%), T604I (13.1%), S711R (7.2%) |
|  | rVSV/SARS-CoV-2<sub>2E1</sub> | 200 | 6 | S254Y (92.1%), N801D (91.5%) |
| RU11 | rVSV/SARS-CoV-2<sub>1D7</sub> | 100 | 6 | I794T (29.3%), N969K (11.6%), K147R (10.3%) |
|  | rVSV/SARS-CoV-2<sub>2E1</sub> | 100 | 6 | C136Y (52.5%), L141P (46.2%), V1128A (46.2%), R682K (6.1%) |
| RU12 | rVSV/SARS-CoV-2<sub>1D7</sub> | 50 | 4 | V143A (6.9%), P1213S (6.5%) |
|  | rVSV/SARS-CoV-2<sub>1D7</sub> | 100 | 4 | E180K (9.3%), V143A (8.3%), H655Y (7.2%), P1213S (5.1%) |
|  | rVSV/SARS-CoV-2<sub>1D7</sub> | 50 | 6 | S813I (7%) |
|  | rVSV/SARS-CoV-2<sub>2E1</sub> | 100 | 4 | L244P (21.3%), T604I (16.6%), N969K (8.7%), M740I (5.6%) |
|  | rVSV/SARS-CoV-2<sub>2E1</sub> | 200 | 6 | N801D (98.9%), S254Y (96.1%), Y144C (5.3%) |
| RU13 | rVSV/SARS-CoV-2<sub>1D7</sub> | 50 | 4 | M153T (14.8%), I794T (10.2%), P1213S (6.5%) |
|  | rVSV/SARS-CoV-2<sub>2E1</sub> | 50 | 3 | S247P (15.9%), N801D (14.5%), T604I (5.5%), P1213S (5.4%) |
|  | rVSV/SARS-CoV-2<sub>2E1</sub> | 50 | 4 | N801D (42.4%), S247P (41.3%), T604I (6.8%), P1213S (6.2%) |
| RU14 | rVSV/SARS-CoV-2<sub>1D7</sub> | 200 | 6 | I794T (6.8%), L212R (5.4%) |
|  | rVSV/SARS-CoV-2<sub>2E1</sub> | 400 | 6 | S399Y (44.9%), T604I (43.6%) |
| RU15 | rVSV/SARS-CoV-2<sub>2E1</sub> | 100 | 4 | T604I (9.3%) |
| RU16 | rVSV/SARS-CoV-2<sub>1D7</sub> | 100 | 2 | N450D (35.8%), V445A (11.9%), K378R (11.8%), V445E (7.8%) |
|  | rVSV/SARS-CoV-2<sub>1D7</sub> | 100 | 6 | V445A (6.8%), G446E (6.1%) |
|  | rVSV/SARS-CoV-2<sub>2E1</sub> | 50 | 5 | G446R (67%), K444R (19%), K444T (9%) |
|  | rVSV/SARS-CoV-2<sub>2E1</sub> | 50 | 6 | V445E (100%), T604I (100%) |
| RU17 | rVSV/SARS-CoV-2<sub>1D7</sub> | 100 | 2 | Q690H (7.8%) |
|  | rVSV/SARS-CoV-2<sub>1D7</sub> | 100 | 4 | Q690H (27.8%) |
|  | rVSV/SARS-CoV-2<sub>1D7</sub> | 100 | 6 | I794T (7.2%) |
|  | rVSV/SARS-CoV-2<sub>2E1</sub> | 100 | 2 | F797S (50.8%), L212R (47%), Q14H (19.6%), T604I (8.4%) |
|  | rVSV/SARS-CoV-2<sub>2E1</sub> | 100 | 4 | Q14H (50%), F797S (37.4%), L212R (36.2%) |
|  | rVSV/SARS-CoV-2<sub>2E1</sub> | 200 | 6 | R102G (33.7%), T604I (33.3%), N87K (29%), H245N (27.2%), P792R (18.5%), I931M (8.3%), F140S (6.3%), R246M (5.4%) |
| RU18 | rVSV/SARS-CoV-2<sub>1D7</sub> | 100 | 2 | R246G (38%), Y248H (35.2%) |
|  | rVSV/SARS-CoV-2<sub>1D7</sub> | 50 | 4 | R246G (55.2%), Y248H (44.5%) |
|  | rVSV/SARS-CoV-2<sub>1D7</sub> | 100 | 4 | R246G (56.9%), Y248H (43.1%) |
|  | rVSV/SARS-CoV-2<sub>1D7</sub> | 100 | 6 | Y248H (26.7%), L176P (9.4%), R682G (8.6%), Y144H (6.3%) |
|  | rVSV/SARS-CoV-2<sub>2E1</sub> | 50 | 4 | P1162S (46.1%), L242P (39.1%), K147N (36.4%), S813I (6.6%) |
|  | rVSV/SARS-CoV-2<sub>2E1</sub> | 100 | 4 | K147N (54.1%), L242P (30%), P1162S (27%) |
|  | rVSV/SARS-CoV-2<sub>2E1</sub> | 200 | 6 | Y144D (32.2%), L244P (27.3%), T604I (18.8%), P174S (16.4%), C136Y (13.9%), L141R (12.2%), G142R (11.2%), V687A (9%), N764S (8.6%), S813I (7%), G261R (5.5%) |
| RU19 | rVSV/SARS-CoV-2<sub>1D7</sub> | 100 | 2 | I794T (11.8%), K150T (6.7%), I1013V (6%) |
|  | rVSV/SARS-CoV-2<sub>1D7</sub> | 100 | 4 | I794T (44%), K150T (13.8%), Y837H (6%) |
|  | rVSV/SARS-CoV-2<sub>1D7</sub> | 100 | 6 | I794T (6.4%) |
|  | rVSV/SARS-CoV-2<sub>2E1</sub> | 200 | 6 | N801D (83.2%), S254Y (79.1%), G142E (13.2%), T573I (6.1%) |
| RU20 | rVSV/SARS-CoV-2<sub>1D7</sub> | 100 | 6 | I794T (8.9%), K147R (7.6%) |
|  | rVSV/SARS-CoV-2<sub>2E1</sub> | 200 | 6 | T604I (82.9%), P25L (60.2%), G142E (30.9%), H69N (7.2%) |
| RU21 | rVSV/SARS-CoV-2<sub>1D7</sub> | 100 | 2 | Y248H (41.4%) |
|  | rVSV/SARS-CoV-2<sub>1D7</sub> | 50 | 4 | Y248H (81.2%) |
|  | rVSV/SARS-CoV-2<sub>1D7</sub> | 100 | 4 | Y248H (87.8%), G769S (5.5%) |
|  | rVSV/SARS-CoV-2<sub>2E1</sub> | 100 | 4 | C15G (74.1%) |
|  | rVSV/SARS-CoV-2<sub>2E1</sub> | 100 | 6 | A243D (95.3%), R190M (6.5%), L244P (5.3%) |
| RU22 | rVSV/SARS-CoV-2<sub>1D7</sub> | 100 | 4 | V1065M (8.4%) |
|  | rVSV/SARS-CoV-2<sub>1D7</sub> | 50 | 6 | V1128A (8%) |
|  | rVSV/SARS-CoV-2<sub>2E1</sub> | 100 | 4 | D808N (18%), T604I (12.4%) |
|  | rVSV/SARS-CoV-2<sub>2E1</sub> | 200 | 6 | R214S (86.1%) |
| RU23 | rVSV/SARS-CoV-2<sub>1D7</sub> | 100 | 2 | E484K (16.6%) |
|  | rVSV/SARS-CoV-2<sub>1D7</sub> | 50 | 4 | E484K (54.1%) |
|  | rVSV/SARS-CoV-2<sub>1D7</sub> | 100 | 4 | T604I (8.4%) |
|  | rVSV/SARS-CoV-2<sub>1D7</sub> | 50 | 6 | I794T (16.4%), Y248S (8.6%), T778N (5.9%) |
|  | rVSV/SARS-CoV-2<sub>2E1</sub> | 100 | 4 | E484K (81.5%) |
| RU24 | rVSV/SARS-CoV-2<sub>1D7</sub> | 100 | 2 | Q493K (10.4%) |
|  | rVSV/SARS-CoV-2<sub>1D7</sub> | 100 | 4 | Y144C (24.7%), T604I (24.6%), I931M (19.9%), E154K (18%) |
|  | rVSV/SARS-CoV-2<sub>1D7</sub> | 100 | 6 | K147E (23.1%), L841Q (18.6%), K147R (14.4%) |
|  | rVSV/SARS-CoV-2<sub>2E1</sub> | 100 | 4 | Q493K (40.5%), L18M (35.6%), N717D (8.3%), L226Q (6%) |
|  | rVSV/SARS-CoV-2<sub>2E1</sub> | 100 | 6 | T604I (100%), Y144C (98.9%) |
| RU25 | rVSV/SARS-CoV-2<sub>1D7</sub> | 50 | 4 | E1092Y (5.1%), R1091S (5.1%) |
|  | rVSV/SARS-CoV-2<sub>1D7</sub> | 100 | 4 | R246G (5.6%) |
|  | rVSV/SARS-CoV-2<sub>2E1</sub> | 100 | 4 | C15Y (27%), P1213S (13.5%), E1188D (8.6%), W1212R (7.7%), K1211R (6.5%), I1210N (5.6%) |
|  | rVSV/SARS-CoV-2<sub>2E1</sub> | 200 | 6 | P25L (65.3%), T604I (14.4%), L244P (10.7%), T734I (6.5%) |
| RU26 | rVSV/SARS-CoV-2<sub>1D7</sub> | 100 | 6 | A623T (78.9%), P681S (77.4%), G142E (54.2%), F759S (19.9%), I931M (12.3%), K113R (9.2%), T778N (5.7%) |
|  | rVSV/SARS-CoV-2<sub>2E1</sub> | 200 | 6 | Y144C (71.7%), T604I (70.5%), I931M (30.6%), F759S (27.7%), F140S (26.4%) |
| RU27 | rVSV/SARS-CoV-2<sub>1D7</sub> | 100 | 6 | K150N (51.5%), K147E (24.1%), L841Q (10.8%), K147R (9%) |
|  | rVSV/SARS-CoV-2<sub>2E1</sub> | 200 | 6 | L244P (79.3%), W64R (75.1%) |
| No plasma | rVSV/SARS-CoV-2<sub>1D7</sub> | 0 | 4 | S98R (31.7%) |
|  | rVSV/SARS-CoV-2<sub>1D7</sub> | 0 | 6 | T604I (73.4%) |
|  | rVSV/SARS-CoV-2<sub>2E1</sub> | 0 | 4 | W64R (31.6%), T604I (13.5%), H66R (7.5%) |
|  | rVSV/SARS-CoV-2<sub>2E1</sub> | 0 | 6 | T604I (28.3%), H66R (17.4%), I931M (8.6%), S982R (8.4%), F759S (7.7%), W64R (6%) |

